# LRRC37A is essential for acrosomal ion homeostasis and male fertility

**DOI:** 10.64898/2026.09.11.751093

**Authors:** Bingbing Wu, Liying Wang, Jiayi Liu, Chenghong Long, Yanjie Ma, Tingting Han, Taicong Tan, Xiaoming Huang, Qingyuan Sun, Wei Li

**Affiliations:** Guangzhou Women and Children’s Medical Center, Guangzhou Medical University, Guangzhou, 510623, China; The Key Laboratory of Organ Regeneration and Reconstruction, State Key Laboratory of Stem Cell and Reproductive Biology, Institute of Zoology, Chinese Academy of Sciences, Beijing, 100049, China; Guangzhou Key Laboratory of Metabolic Diseases and Reproductive Health, Guangdong-Hong Kong Metabolism & Reproduction Joint Laboratory, Reproductive Medicine Center, The Affiliated Guangdong Second Provincial General Hospital of Jinan University, Guangzhou, 510317, China; Key Laboratory of Regenerative Medicine of Ministry of Education, Jinan University, Guangzhou, 510632, China; Department of Developmental Biology, School of Basic Medical Sciences, Southern Medical University, Guangzhou, 510515, China

**Keywords:** Acrosome, LRRC37A, Na/K-ATPase, Ion homeostasis, Male infertility

## Abstract

Mammalian spermatozoa face various challenges during their long march to the destination. As a specialized membrane-bound organelle in the sperm head, the acrosome is particularly sensitive to changes in ionic and osmotic conditions, yet how acrosomal ion homeostasis is maintained remains largely unknown. Here, we show that LRRC37A is required for acrosomal ion homeostasis. Loss of *Lrrc37a* causes complete male infertility, accompanied by pronounced enlargement and deformation of the acrosome in the sperm head. Acrosome biogenesis was not affected by *Lrrc37a* knockout during the early stages of spermiogenesis, whereas acrosomal enlargement emerged at later stages and became more pronounced in the epididymis. LRRC37A localized to the outer acrosomal region and interacted with the Na/K-ATPase α-subunit isoforms ATP1A1 and ATP1A4. Loss of LRRC37A disrupted the spatial distribution of both isoforms and was accompanied by increased Na⁺ within the acrosomal region. In addition, pharmacological perturbation of Na/K-ATPase activity or cellular ion homeostasis induced acrosomal swelling. These findings establish an essential role for LRRC37A in acrosomal ion homeostasis, suggesting that its deficiency may be associated with a distinct form of teratozoospermia characterized by acrosomal enlargement.

## Introduction

Mammalian spermatozoa face various challenges from changing extracellular environments as they move from the testis to the epididymis and ultimately into the female reproductive tract. Along this journey, they are exposed to marked variations in pH, viscosity, fluid dynamics, ionic composition, and osmolality ^1,2^. The epididymal lumen is relatively hyperosmotic, whereas the uterus and oviduct exhibit markedly different osmotic and ionic conditions ^1,3^. Successful maturation and fertilization therefore require sperm to adapt rapidly to substantial physicochemical changes while preserving functional competence ^4,5^.

As highly differentiated cells with limited capacity for *de novo* gene expression, mature spermatozoa rely predominantly on pre-existing proteins to respond to extracellular cues ^6–9^. Membrane ion channels and transporters are central to this process. Dynamic regulation of Na⁺, K⁺, Ca²⁺, H⁺, and HCO₃⁻ contributes to the control of capacitation, hyperactivated motility, and acrosomal exocytosis ^4^. Ion transport is also tightly coupled to osmotic balance and cell-volume regulation, enabling sperm to accommodate the osmotic challenges encountered during epididymal maturation and transit through the female reproductive tract ^2,3^.

At the same time, sperm must preserve the structural and functional integrity of highly specialized membrane domains. Fertilization involves a sequence of membrane-dependent events, including interactions with the cumulus and zona pellucida (ZP), acrosomal exocytosis, and extensive remodeling of sperm-head membranes ^5,10,11^. These processes require sperm-head membranes to remain structurally stable during maturation while retaining the capacity for rapid reorganization at fertilization. Ionic and osmotic balance is closely linked to sperm volume regulation and membrane integrity ^2,12^. As a specialized membrane-bound organelle in the sperm head, the acrosome maintains a distinct luminal environment and is highly sensitive to osmotic perturbation ^13^, raising the possibility that local ion homeostasis contributes to the preservation of acrosomal membrane organization and morphology.

The acrosome is a Golgi-derived membrane-bound organelle that develops over the anterior surface of the sperm nucleus during spermiogenesis ^14,15^. Its formation involves the trafficking and fusion of Golgi-derived proacrosomal vesicles, followed by spreading and remodeling of the resulting acrosomal vesicle over the anterior nuclear surface. In mature sperm, the acrosomal compartment is bounded by the outer acrosomal membrane (OAM) and inner acrosomal membrane (IAM), with the sperm plasma membrane closely apposed to the OAM ^14,15^. Most studies of acrosomal biology have focused either on its biogenesis during spermiogenesis or on its exocytotic remodeling during fertilization ^15–17^. Less is known about how the established acrosomal compartment is maintained during the interval between these two stages. During late spermiogenesis and epididymal maturation, the acrosome must preserve its morphology and internal environment before undergoing tightly regulated exocytosis at fertilization. Despite the established importance of ion transport in sperm physiology, how ionic homeostasis is maintained within the acrosomal region remains unclear.

LRRC37A belongs to the leucine-rich repeat-containing 37 (*LRRC37*) gene family. Comparative expression analyses indicate that testis-restricted expression likely represents the ancestral state of the *LRRC37* family ^18^. In mice, the two *Lrrc37* genes are expressed only in the testis, whereas macaque and human *LRRC37* genes show broader expression profiles while retaining prominent testicular expression. LRRC37 proteins are predicted to be single-span transmembrane proteins, and heterologous expression studies showed that recombinant human LRRC37A traffics through the Golgi apparatus to the plasma membrane ^18^. However, the physiological role of LRRC37A in male reproduction, particularly whether it contributes to acrosomal function, remains unknown.

Here, we define a role for LRRC37A in acrosomal ion homeostasis. *Lrrc37a* knockout in mice leads to complete male infertility, accompanied by impaired acrosomal function and pronounced acrosomal enlargement. Acrosomal swelling emerged during the late stages of spermiogenesis and became more pronounced in epididymal spermatozoa. Functionally, LRRC37A was positioned toward the outer acrosomal region and interacted with the Na/K-ATPase α-subunit isoforms ATP1A1 and ATP1A4. Loss of LRRC37A disrupted the spatial distribution of both isoforms and was accompanied by increased Na⁺ within the acrosomal region. These findings reveal an essential role for LRRC37A in acrosomal ion homeostasis and link local ionic regulation in the sperm head to normal acrosomal function, suggesting that LRRC37A deficiency may be associated with a distinct form of teratozoospermia characterized by acrosomal enlargement.

## Results

### LRRC37A is a conserved sperm-head protein required for male fertility

To investigate the potential function of LRRC37A in male reproduction, we first examined the evolutionary distribution and reproductive expression profile of LRRC37A. Phylogenetic analysis revealed that LRRC37A is conserved across diverse internally fertilizing vertebrates, including cartilaginous fishes, reptiles, birds, and mammals (Fig. 1A). In mice, *Lrrc37a* expression was strongly enriched in the testis and increased during postnatal testicular development (Fig. 1B, C). Immunofluorescence further showed that LRRC37A was restricted to the anterior sperm head, with a similar localization pattern observed in human spermatozoa (Fig. 1D). Together, these observations suggested a conserved reproductive role for LRRC37A associated with the sperm head.

**Figure 1.**
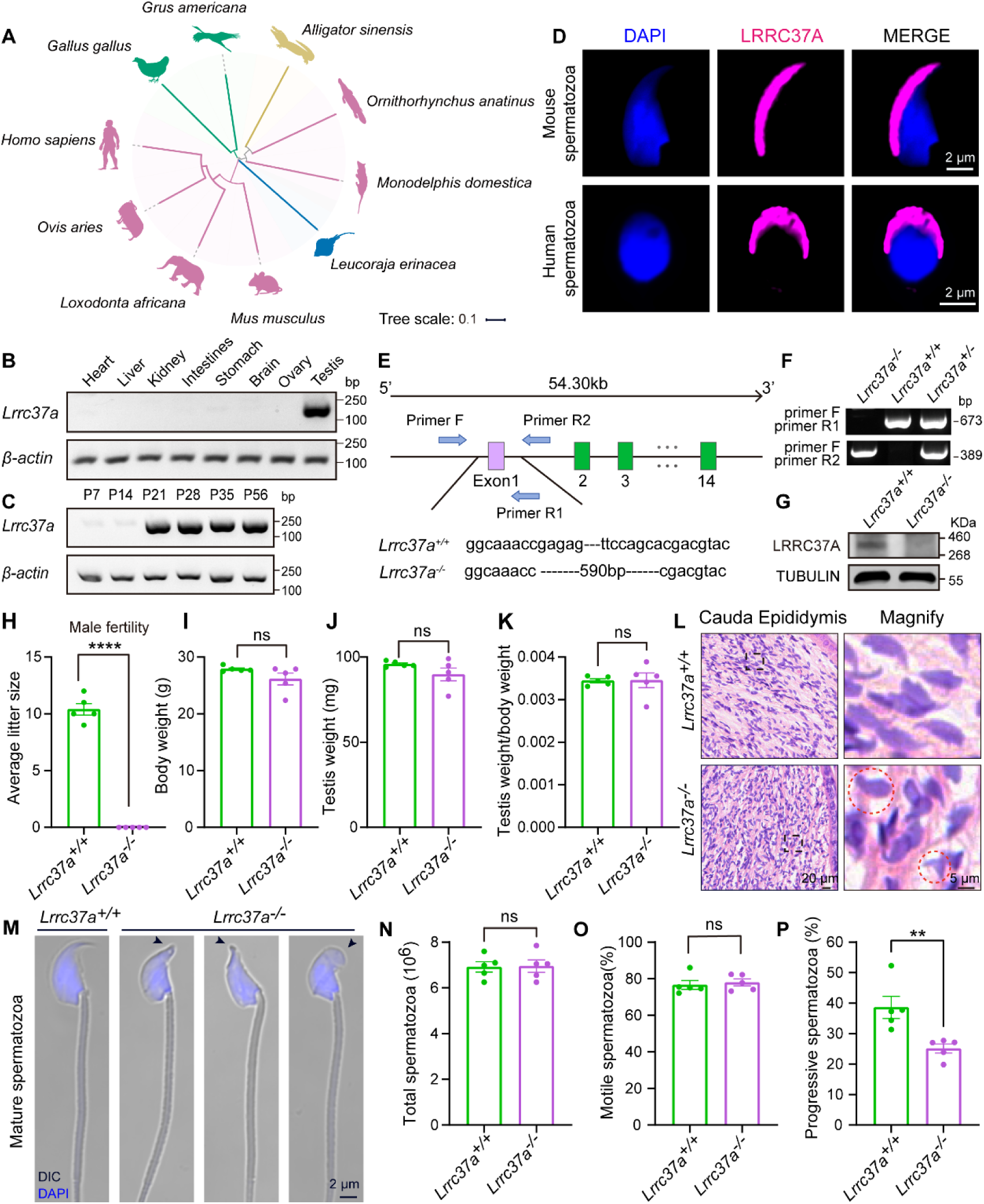
LRRC37A localizes to the sperm head and is essential for male fertility. (A) Phylogenetic analysis of LRRC37A homologues in representative internally fertilizing vertebrates. Colors indicate major vertebrate groups: birds (green), reptiles (yellow), mammals (pink), and cartilaginous fishes (blue). Tree scale indicates the number of amino acid substitutions per site. (B) Reverse transcription polymerase chain reaction (RT-PCR) analysis of *Lrrc37a* expression in adult mouse tissues. *β-actin* served as the loading control. (C) RT-PCR analysis of *Lrrc37a* expression in testes collected at the indicated postnatal ages. *β-actin* served as the loading control. (D) Immunofluorescence localization of LRRC37A in mouse and human spermatozoa. DNA was counterstained with DAPI. (E) Schematic of the *Lrrc37a* locus showing the 590 bp deletion introduced into exon 1 and the positions of the genotyping primers. (F) Representative PCR genotyping of *Lrrc37a*^+/+^, *Lrrc37a*^+/–^, and *Lrrc37a*^−/−^ mice. (G) Immunoblot analysis confirming loss of LRRC37A protein in *Lrrc37a*^−/−^ testes. TUBULIN was used as the loading control. (H) Average litter size obtained from fertility tests of *Lrrc37a*^+/+^ and *Lrrc37a*^−/−^ male. Data are presented as the mean ± SEM. Two-tailed, unpaired Student’s t test: \*\*\*\**P* < 0.0001, *n* = 5 independent experiments. (I–K) Body weight (I), testis weight (J), and testis-to-body weight ratio (K) of adult *Lrrc37a*^+/+^ and *Lrrc37a*^−/−^ males. Data are presented as the mean ± SEM. Two-tailed, unpaired Student’s t test: ns, not significant, *n* = 5 independent experiments. (L) Hematoxylin and eosin staining of cauda epididymides from *Lrrc37a*^+/+^ and *Lrrc37a*^−/−^ males. Red dashed circles indicate spermatozoa with abnormal head morphology within the epididymal lumen. (M) Representative DIC images of spermatozoa released from the cauda epididymis. DNA was stained with DAPI. (N–P) Total sperm number in the cauda epididymis (N), percentage of total motile spermatozoa (O), and percentage of progressively motile spermatozoa (P). Data are presented as the mean ± SEM. Two-tailed, unpaired Student’s t test: \*\**P* < 0.01; ns, not significant, *n* = 5 independent experiments.

To determine whether LRRC37A is required for male fertility, we generated *Lrrc37a* deficient mice carrying a 590 bp deletion in exon 1 and confirmed the loss of LRRC37A protein in homozygous testes (Fig. 1E–G). Consistent with this, the characteristic LRRC37A signal in the head of mature spermatozoa was also absent in *Lrrc37a*^−/−^ males (Fig. S1A). *Lrrc37a*^−/−^ males were infertile (Fig. 1H), whereas body weight, testis weight, and the testis-to-body weight ratio were comparable with those of control males (Fig. 1I–K). Histological examination of the cauda epididymis revealed spermatozoa with abnormal head morphology within the epididymal lumen of *Lrrc37a*^−/−^ males (Fig. 1L, red dashed circles). Consistent with this observation, spermatozoa released from the cauda epididymis of *Lrrc37a*^−/−^ males displayed pronounced head shape abnormalities, characterized by blunting and enlargement of the anterior sperm head (Fig. 1M, black arrowheads).

Total sperm numbers in the cauda epididymis were comparable between *Lrrc37a*^+/+^ and *Lrrc37a*^−/−^ males (Fig. 1N), and total sperm motility did not differ significantly between the two groups (Fig. 1O). In contrast, progressive motility was significantly reduced in *Lrrc37a*^−/−^ spermatozoa (Fig. 1P). Together, these findings establish LRRC37A as a sperm-head protein required for male fertility, with its loss resulting in pronounced sperm-head abnormalities and reduced progressive motility.

### LRRC37A deficiency leads to complete fertilization failure due to impaired acrosome function

We next directly assessed the fertilization competence of *Lrrc37a*^−/−^ spermatozoa (Fig. 2A). In conventional *in vitro* fertilization (IVF) assay using cumulus-intact oocytes, *Lrrc37a*^+/+^ spermatozoa efficiently supported fertilization, with 93.44 ± 1.12% of oocytes forming two pronuclei at 6 h post-insemination. In contrast, no two-pronuclear oocytes were detected following insemination with *Lrrc37a*^−/−^ spermatozoa, and the fertilization rate was 0% (Fig. 2B–D).

**Figure 2.**
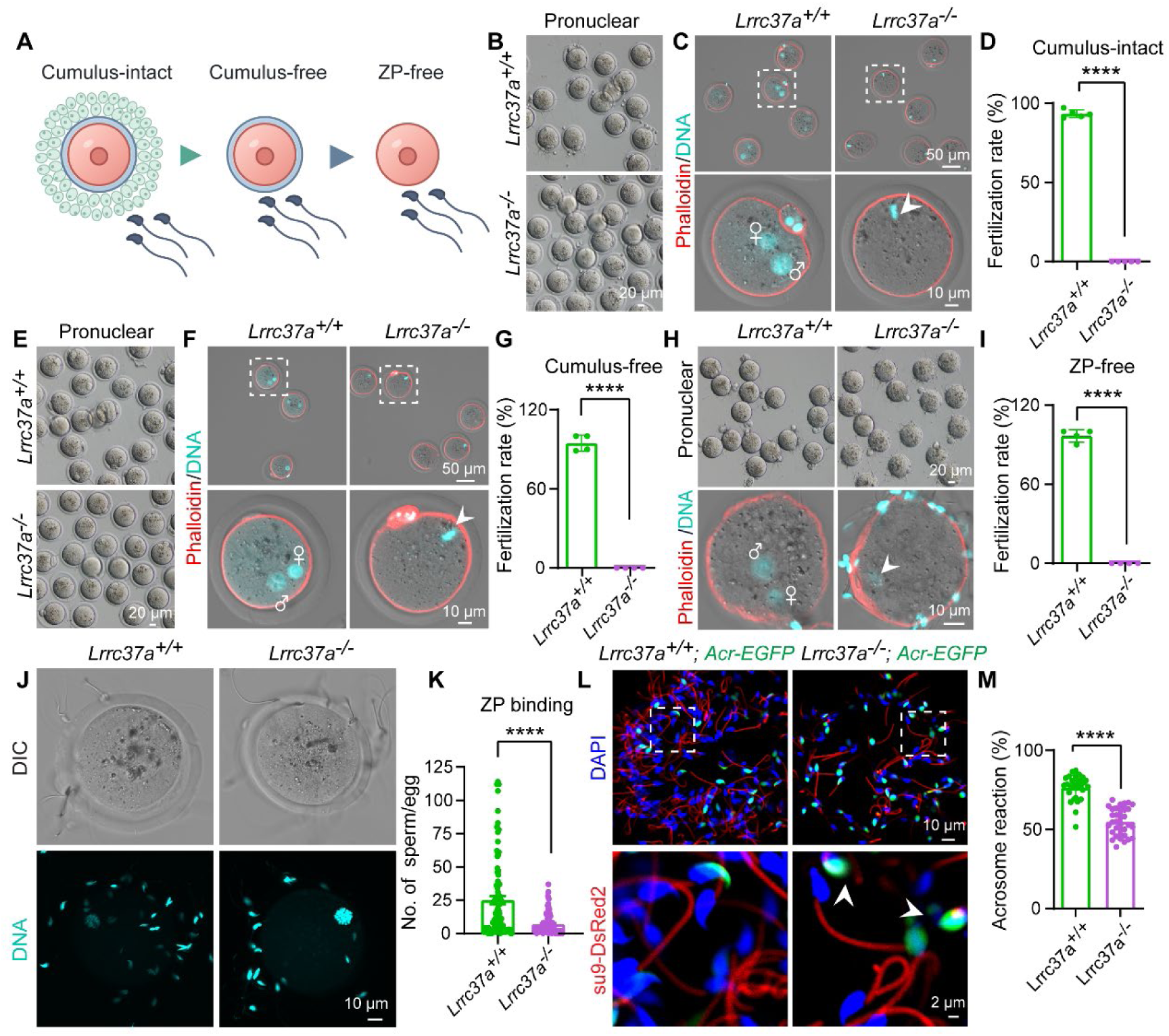
*Lrrc37a* deficient spermatozoa fail to fertilize oocytes due to impaired acrosome function. (A) Schematic of the *in vitro* fertilization (IVF) assays performed with cumulus-intact, cumulus-free, and zona pellucida (ZP)-free oocytes. (B) The fertilizing ability of *Lrrc37a*^+/+^ and *Lrrc37a*^−/−^ spermatozoa was evaluated using cumulus-intact oocytes in an IVF assay at 6 hours post-insemination. (C) Representative fluorescence images of oocytes from the assay in (B). F-actin was labeled with Phalloidin (red), and DNA was stained with DAPI (cyan). Fertilized oocytes exhibiting two pronuclei (2PN) were observed in the *Lrrc37a*^+/+^ spermatozoa group, whereas no 2PN oocytes were detected following insemination with *Lrrc37a*^−/−^ spermatozoa. Female and male pronuclei are indicated by ♀ and ♂, respectively. The white arrowhead indicates chromosomes in unfertilized oocytes. (D) Fertilization rates of cumulus-intact oocytes inseminated with *Lrrc37a*^+/+^ and *Lrrc37a*^−/−^ spermatozoa from the assay in (B). Fertilization rates: *Lrrc37a*^+/+^, 93.44 ± 1.12% (total oocytes examined = 177); *Lrrc37a*^−/−^, 0.00 ± 0.00%, (total oocytes examined = 233). Data are presented as the mean ± SEM. Two-tailed, unpaired Student’s t test: \*\*\*\**P* < 0.0001, *n* = 3 independent experiments. (E) The fertilizing ability of *Lrrc37a*^+/+^ and *Lrrc37a*^−/−^ spermatozoa was evaluated using cumulus-free oocytes in an IVF assay at 6 hours post-insemination. (F) Representative fluorescence images of oocytes from the assay in (E). F-actin was labeled with Phalloidin (red), and DNA was stained with DAPI (cyan). Fertilized eggs exhibiting two pronuclei (2PN) were observed in the *Lrrc37a*^+/+^ spermatozoa group, whereas no 2PN oocytes were detected following insemination with *Lrrc37a*^−/−^ spermatozoa. Female and male pronuclei are indicated by ♀ and ♂, respectively. The white arrowhead indicates chromosomes in unfertilized oocytes. (G) Fertilization rates of cumulus-free oocytes inseminated with *Lrrc37a*^+/+^ and *Lrrc37a*^−/−^ spermatozoa from the assay in (E). Fertilization rates: *Lrrc37a*^+/+^, 94.73 ± 3.05% (total oocytes examined = 86); *Lrrc37a*^−/−^, 0.00 ± 0.00%, total oocytes examined = 191). Data are presented as the mean ± SEM. Two-tailed, unpaired Student’s t test: \*\*\*\**P* < 0.0001, *n* = 3 independent experiments. (H) The fertilizing ability of *Lrrc37a*^+/+^ and *Lrrc37a*^−/−^ spermatozoa was evaluated using ZP-free oocytes in an IVF assay at 6 hours post-insemination. Representative fluorescence images of pronuclei stained with Phalloidin (red) and DAPI (cyan). Fertilized eggs exhibiting two pronuclei (2PN) were observed in the *Lrrc37a*^+/+^ spermatozoa group, whereas no 2PN oocytes were detected following insemination with *Lrrc37a*^−/−^ spermatozoa. The sex of the pronuclei is indicated with ♀ and ♂ symbols, and white arrowhead indicates chromosomes within unfertilized oocytes. (I) Fertilization rates of ZP-free oocytes inseminated with *Lrrc37a*^+/+^ and *Lrrc37a*^−/−^ spermatozoa from the assay in (H). Fertilization rates: *Lrrc37a*^+/+^, 96.73 ± 2.35% (total oocytes examined = 118); *Lrrc37a*^−/−^, 0.00 ± 0.00%, (total oocytes examined = 175). Data are presented as the mean ± SEM. Two-tailed, unpaired Student’s t test: \*\*\*\**P* < 0.0001, *n* = 3 independent experiments. (J) Zona pellucida (ZP) binding assay. Wild-type MII oocytes were incubated with capacitated *Lrrc37a*^+/+^ or *Lrrc37a*^−/−^ spermatozoa for 1 hour. DNA was stained with DAPI (cyan). (K) Quantification of spermatozoa bound to the ZP surface per oocyte. *Lrrc37a*^+/+^, 25.05 ± 3.01 (total oocytes examined = 95); *Lrrc37a*^−/−*-*^, 6.88 ± 0.76 (total oocytes examined = 100). Data are presented as the mean ± SEM. Two-tailed, unpaired Student’s t test: \*\*\*\**P* < 0.0001. (L) Acrosome reaction assay under fertilization conditions using *Lrrc37a*^+/+^ and *Lrrc37a*^−/−^ spermatozoa carrying a transgene expressing EGFP in the acrosome and DsRed2 in the mitochondria. Capacitated spermatozoa were co-incubated with wild-type MII oocytes for 1.5 h, allowing the acrosome reaction to be assessed in the context of sperm–oocyte interaction. Loss of acrosomal EGFP fluorescence was used to identify acrosome-reacted spermatozoa. White arrowheads indicate spermatozoa with swollen acrosomes. (M) Quantification of acrosome reaction rates from the assay in (L). *Lrrc37a*^+/+^, 76.39 ± 1.40 % (total oocytes examined = 30); *Lrrc37a*^−/−^, 54.92 ± 1.57 % (total oocytes examined = 31). Data are presented as the mean ± SEM. Two-tailed, unpaired Student’s t test: \*\*\*\**P* < 0.0001.

To determine whether the fertilization failure resulted from an inability to traverse the cumulus matrix, IVF was performed using cumulus-free oocytes. Removal of the cumulus did not restore fertilization by *Lrrc37a*^−/−^ spermatozoa. Whereas 94.73 ± 3.05% of oocytes were fertilized by control spermatozoa, no two-pronuclear oocytes were detected in the knockout group (Fig. 2E–G). The zona pellucida was then removed to bypass an additional extracellular barrier. *Lrrc37a*^+/+^ spermatozoa fertilized 96.73 ± 2.35% of ZP-free oocytes, whereas fertilization by *Lrrc37a*^−/−^ spermatozoa remained completely absent (Fig. 2H, I). Thus, the complete fertilization failure of *Lrrc37a*^−/−^ spermatozoa cannot be attributed solely to impaired penetration of the cumulus matrix or zona pellucida.

Sperm functions related to the acrosomal region were then assessed. In the zona pellucida-binding assay, the number of sperm bound per oocyte was markedly reduced in the knockout group, from 25.05 ± 3.01 in controls to 6.88 ± 0.76 in *Lrrc37a*^−/−^ spermatozoa (Fig. 2J, K). To assess the acrosome reaction during sperm–oocyte interaction, we crossed *Lrrc37a*^−/−^ mice with transgenic mice expressing EGFP in the acrosome and DsRed2 in the mitochondria ^19^, allowing the acrosomal status of individual spermatozoa to be detected by fluorescence microscopy. Capacitated spermatozoa were then co-incubated with MII oocytes, and loss of the acrosomal EGFP signal was used to identify acrosome-reacted spermatozoa. The proportion of acrosome-reacted spermatozoa was significantly lower in the *Lrrc37a*^−/−^ group than in controls (54.92 ± 1.57% versus 76.39 ± 1.40%; Fig. 2L, M). Together, these findings show that LRRC37A deficiency causes complete fertilization failure under cumulus-intact, cumulus-free, and ZP-free conditions and is accompanied by impaired zona pellucida binding and a reduced acrosome reaction.

### LRRC37A deficiency causes pronounced acrosomal swelling in epididymal spermatozoa

During the acrosome reaction assay, we noticed that many *Lrrc37a*^−/−^ spermatozoa exhibited an enlarged acrosomal region (Fig.2L, white arrowheads). Freshly released live spermatozoa from the cauda epididymis were analyzed using the acrosomal EGFP reporter, confirming pronounced enlargement of the acrosomal compartment in *Lrrc37a*^−/−^ spermatozoa. To determine whether this abnormality developed during epididymal transit, spermatozoa from the corpus and caput epididymides were also analyzed. Similar acrosomal enlargement was already evident in both regions, indicating that the defect was present before sperm reached the cauda epididymis and persisted during epididymal maturation (Fig. 3A). Scanning electron microscopy further revealed marked abnormalities at the anterior sperm head. Compared with the compact and well-defined acrosomal region of control spermatozoa, *Lrrc37a*^−/−^ spermatozoa exhibited pronounced expansion and deformation of the acrosomal region (Fig. 3B, white arrowheads).

**Figure 3.**
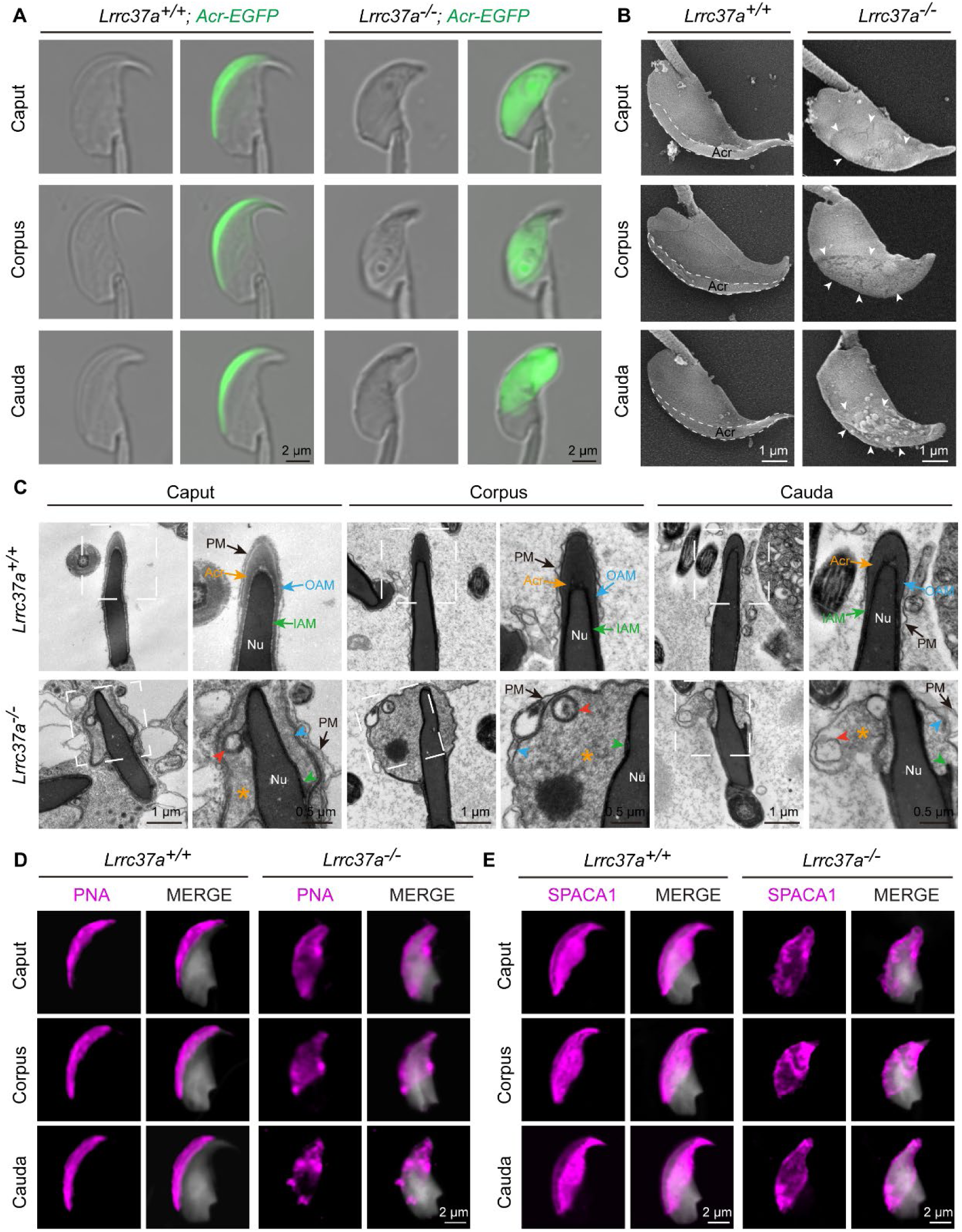
LRRC37A deficiency causes acrosomal expansion and disrupts membrane organization. (A) Representative DIC and acrosomal EGFP images of freshly released live spermatozoa isolated from the caput, corpus, and cauda epididymides of *Lrrc37a*^+/+^ and *Lrrc37a*^−/−^ mice. *Lrrc37a*^−/−^ spermatozoa show marked expansion and distortion of the acrosomal region. (B) Scanning electron microscopy of epididymal sperm heads. Dashed lines outline the acrosomal region in control spermatozoa. Arrowheads indicate abnormal expansion and deformation of the acrosomal region in *Lrrc37a*^−/−^ spermatozoa. (C) Transmission electron microscopy of sperm heads isolated from the caput, corpus, and cauda epididymides. Boxed regions are shown at higher magnification in the adjacent panels. In *Lrrc37a*^−/−^ spermatozoa, colored arrowheads indicate disorganization of the plasma and acrosomal membranes. Orange asterisks indicate expansion of the acrosomal compartment, and red arrowheads indicate abnormal vesicular membrane structures. PM, plasma membrane; OAM, outer acrosomal membrane; IAM, inner acrosomal membrane; Acr, acrosome; Nu, nucleus. (D) PNA staining of spermatozoa isolated from the caput, corpus, and cauda epididymides of *Lrrc37a*^+/+^ and *Lrrc37a*^−/−^ mice. DNA was stained with DAPI and is shown in gray. (E) SPACA1 staining of spermatozoa isolated from the caput, corpus, and cauda epididymides of *Lrrc37a*^+/+^ and *Lrrc37a*^−/−^ mice. DNA was stained with DAPI and is shown in gray.

Transmission electron microscopy was then used to examine the organization of the acrosomal region in spermatozoa from different regions of the epididymis. In control spermatozoa, the plasma membrane, outer acrosomal membrane, and inner acrosomal membrane were closely apposed and maintained an ordered spatial arrangement, while the acrosomal compartment showed a compact, electron-dense appearance (Fig. 3C). In *Lrrc37a*^−/−^ spermatozoa, these membranes were irregular and disorganized, accompanied by pronounced expansion of the acrosomal compartment with markedly reduced electron density throughout the epididymis (Fig. 3C, orange asterisks) and the appearance of abnormal vesicular membrane structures (Fig. 3C, red arrowheads).

To further assess the acrosomal abnormalities, the distributions of PNA, a lectin that labels the outer acrosomal membrane ^20^, and SPACA1, an inner acrosomal membrane-associated protein ^21^, were analyzed. Both PNA and SPACA1 showed markedly altered distribution patterns in *Lrrc37a*^−/−^ spermatozoa from the caput, corpus, and cauda epididymides (Fig. 3D, E). These findings demonstrate pronounced expansion of the acrosomal compartment following LRRC37A loss, accompanied by extensive structural abnormalities of the surrounding membrane system.

### Acrosomal abnormalities emerge during late spermiogenesis in *Lrrc37a* deficient spermatids

Given that acrosomal abnormalities were already present in spermatozoa from different regions of the epididymis, we asked when these defects first emerged during sperm development. Testis sections were examined by periodic acid–Schiff (PAS) staining, which was used to identify the 12 stages of the seminiferous epithelial cycle based largely on spermatid and acrosomal morphology ^22^. The overall progression of spermatogenesis was preserved in *Lrrc37a*^−/−^ testes. However, late-stage spermatids present from stages I–III through VII–VIII showed a distinctly blunted anterior head (Fig. 4A, red arrowheads). Acrosome morphology across spermiogenesis was further assessed by PNA staining. No obvious differences were detected between *Lrrc37a*^+/+^ and *Lrrc37a*^−/−^ spermatids during the early stages. In contrast, distortion and enlargement of the anterior acrosomal region became evident in steps 13–16 *Lrrc37a*^−/−^ spermatids (Fig. 4B, white arrowheads).

**Figure 4.**
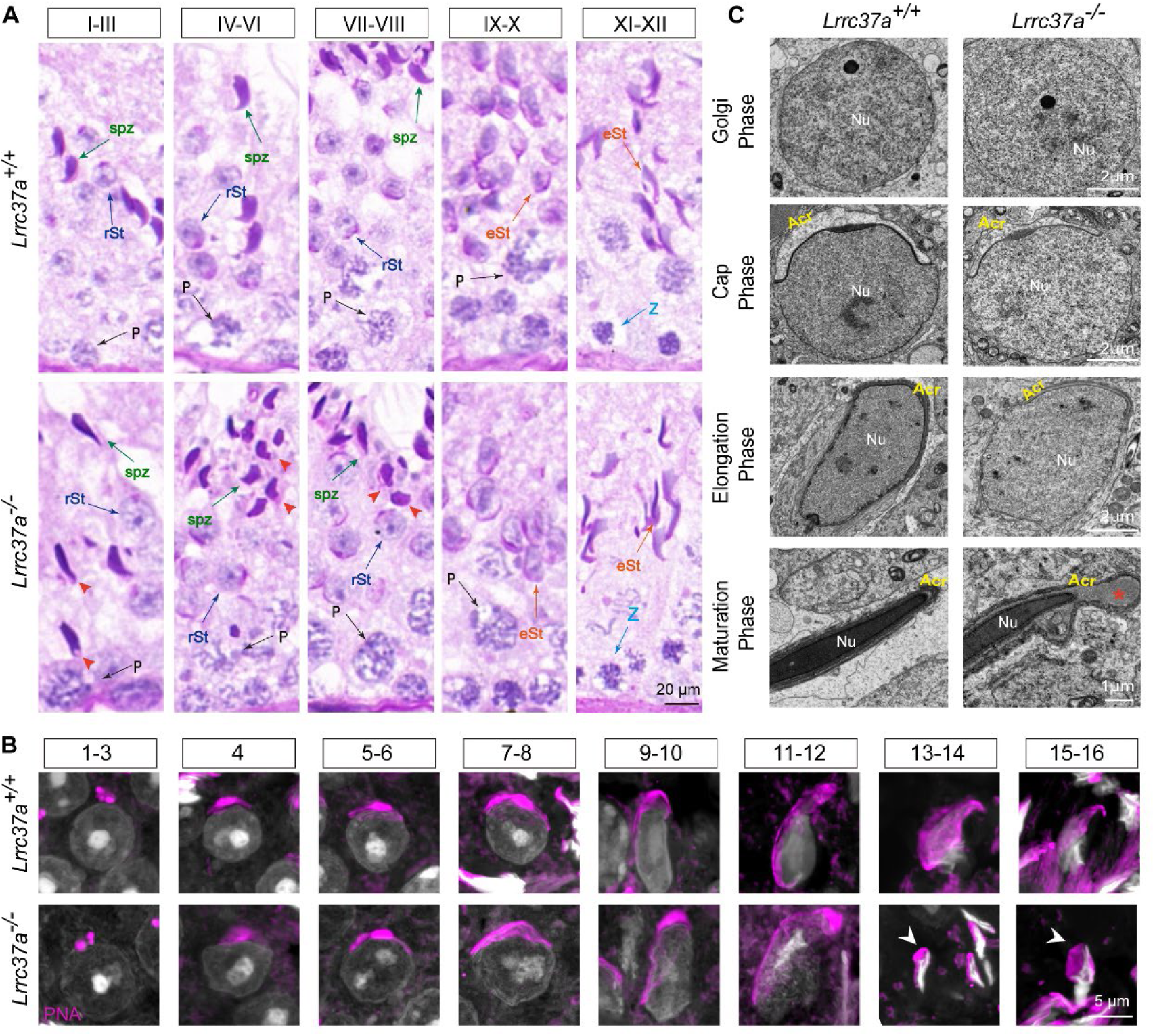
Acrosomal abnormalities emerge during the late stages of spermiogenesis in *Lrrc37a* deficient spermatids. (A) Periodic acid–Schiff (PAS) staining of testis sections from *Lrrc37a*^+/+^ and *Lrrc37a*^−/−^ mice showing representative stages I–XII of the seminiferous epithelial cycle. Red arrowheads indicate blunting of the anterior head in late-stage *Lrrc37a*^−/−^ spermatids present from stages I– III through VII–VIII. spz, spermatozoa; rSt, round spermatids; eSt, elongating spermatids; P, pachytene spermatocytes; Z, zygotene spermatocytes. (B) PNA staining of developing spermatids at representative steps of spermiogenesis in *Lrrc37a*^+/+^ and *Lrrc37a*^−/−^ testes. White arrowheads indicate abnormal acrosomal morphology. DNA was stained with DAPI and is shown in gray. (C) Transmission electron microscopy of spermatids during the Golgi, cap, elongation, and maturation phases of spermiogenesis. Acrosome formation appears normal during the Golgi, cap, and elongation phases in *Lrrc37a*^−/−^ spermatids, whereas marked expansion and structural disorganization of the acrosomal compartment become evident during the maturation phase. Red asterisks indicate abnormal expansion of the acrosomal compartment. Acr, acrosome; Nu, nucleus.

Transmission electron microscopy further defined the developmental onset of the defect. Acrosome formation appeared normal during the Golgi, cap, and elongation phases in *Lrrc37a*^−/−^spermatids. In the maturation phase, the acrosomal compartment became markedly expanded and structurally disorganized (Fig. 4C, red asterisk). Thus, the acrosomal defect caused by LRRC37A deficiency emerges during the late stages of spermiogenesis.

### LRRC37A localizes to the outer acrosomal region and interacts with the Na/K-ATPase α-subunit isoforms ATP1A1 and ATP1A4

To explore the molecular basis of the acrosomal abnormalities caused by LRRC37A deficiency, we first examined its subcellular localization in developing spermatids at different stages of spermiogenesis. LRRC37A remained closely localized to the acrosomal region throughout spermiogenesis (Fig. 5A).

**Figure 5.**
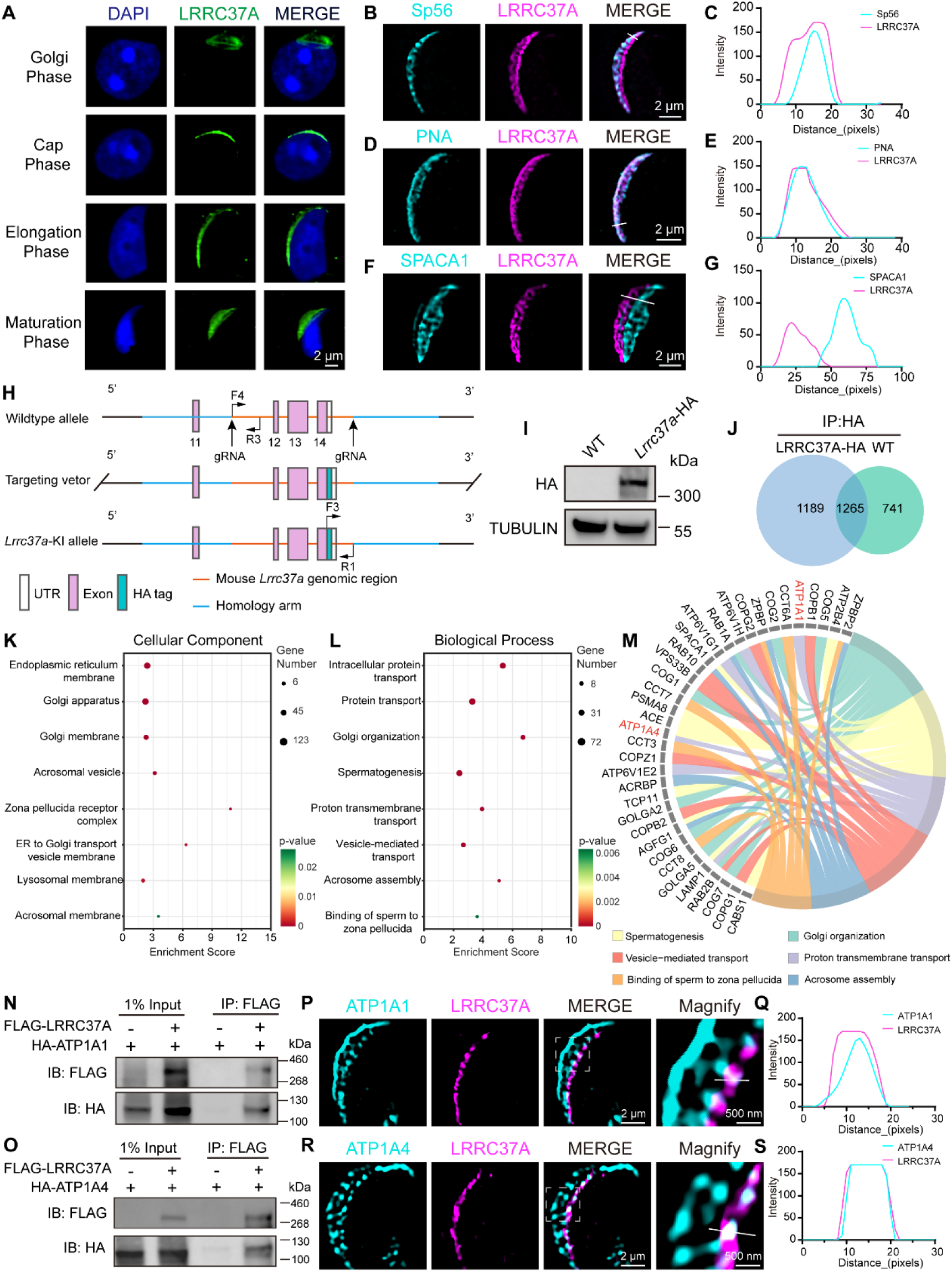
Spatial localization of LRRC37A and its interaction with ATP1A1 and ATP1A4. (A) Immunofluorescence analysis of LRRC37A in developing spermatids at representative stages of spermiogenesis. DNA was stained with DAPI. DNA was counterstained with DAPI. (B, C) Structured illumination microscopy (SIM) analysis of the spatial relationship between LRRC37A and Sp56 in spermatozoa, with corresponding line-scan fluorescence intensity profiles. (D, E) SIM analysis of the spatial relationship between LRRC37A and PNA in spermatozoa, with corresponding line-scan fluorescence intensity profiles. (F, G) SIM analysis of the spatial relationship between LRRC37A and SPACA1 in spermatozoa, with corresponding line-scan fluorescence intensity profiles. (H) Schematic of the strategy used to generate HA-tagged *Lrrc37a* knock-in mice. The positions of the guide RNAs and genotyping primers are indicated. (I) Immunoblot analysis of HA-tagged LRRC37A in testes from HA-tagged *Lrrc37a* knock-in and wild-type mice. TUBULIN served as the loading control. (J) Venn diagram showing proteins identified by anti-HA immunoprecipitation–mass spectrometry in HA-tagged *Lrrc37a* knock-in and wild-type testes. (K, L) Gene Ontology enrichment analysis of candidate LRRC37A-interacting proteins for cellular component (K) and biological process (L). (M) Chord diagram showing relationships between candidate LRRC37A-interacting proteins and enriched biological processes. (N, O) Co-immunoprecipitation analysis of FLAG-LRRC37A with HA-ATP1A1 (N) or HA-ATP1A4 (O) in co-transfected HEK293T cells. (P, Q) SIM analysis of the spatial relationship between LRRC37A and ATP1A1 in the sperm head, with corresponding line-scan fluorescence intensity profiles. Boxed regions are enlarged at right. (R, S) SIM analysis of the spatial relationship between LRRC37A and ATP1A4 in the sperm head, with corresponding line-scan fluorescence intensity profiles. Boxed regions are enlarged at right.

Structured illumination microscopy (SIM) was then used to define the position of LRRC37A within the acrosome at higher spatial resolution. Comparison with Sp56, a well-characterized acrosome-associated protein ^23^, showed partial colocalization between the two signals (Fig. 5B, C). The position of LRRC37A was further compared with PNA, which labels the outer acrosomal membrane, and SPACA1, an inner acrosomal membrane-associated protein. LRRC37A showed almost complete overlap with PNA (Fig. 5D, E), whereas its signal was positioned external to SPACA1 (Fig. 5F, G). Together, these spatial relationships place LRRC37A toward the outer region of the acrosomal compartment.

To identify endogenous LRRC37A-interacting proteins, we generated an HA-tagged *Lrrc37a*knock-in mouse (Fig. 5H). Expression of HA-tagged LRRC37A was confirmed by immunoblotting (Fig. 5I), and immunofluorescence showed a localization pattern comparable to that detected with the LRRC37A antibody. In mature spermatozoa, the HA signal showed extensive overlap with PNA (Fig. S1B, C) and was positioned external to SPACA1 (Fig. S1D, E), further confirming that HA-tagged LRRC37A retained the characteristic acrosomal localization of the endogenous protein. Immunoprecipitation–mass spectrometry (IP–MS) was subsequently performed using testicular lysates from HA-*Lrrc37a* knock-in mice, with wild-type testes included as a negative control and subjected to the same anti-HA immunoprecipitation procedure. A total of 1,189 proteins were detected exclusively in the HA-*Lrrc37a* samples and not in wild-type controls, and were therefore considered candidate LRRC37A-interacting proteins (Fig. 5J, Table S1).

Gene Ontology analysis of these proteins revealed enrichment of membrane-related cellular components, including the Golgi apparatus, Golgi membrane, acrosomal vesicle, and acrosomal membrane, together with biological processes related to acrosome function, vesicle-mediated transport, and transmembrane transport (Fig. 5K–M). Among the candidate proteins, ATP1A1 and ATP1A4 were of particular interest as Na/K-ATPase α-subunit isoforms expressed in the testis and sperm ^24^. ATP1A1 is broadly expressed, whereas ATP1A4 is largely restricted to male germ cells and is essential for sperm ion homeostasis and fertilization ^25^. Recent super-resolution imaging further showed that both isoforms are present in the sperm head and exhibit distinct, highly organized spatial distributions ^26^. Co-immunoprecipitation confirmed interactions of LRRC37A with both ATP1A1 and ATP1A4 (Fig. 5N, O). We next examined their spatial relationship with LRRC37A in the sperm head by structured illumination microscopy. Both ATP1A1 and ATP1A4 were positioned external to LRRC37A, with partial overlap between the signals (Fig. 5P-S). Together, these findings identify ATP1A1 and ATP1A4 as LRRC37A-interacting proteins that occupy closely apposed regions of the sperm head.

### LRRC37A is required for Na/K-ATPase localization and acrosomal Na⁺ homeostasis

We then assessed whether LRRC37A is required to maintain the normal localization of ATP1A1 and ATP1A4 in spermatozoa. In *Lrrc37a*^+/+^ spermatozoa, ATP1A1 displayed a defined distribution along the sperm head and partially overlapped with the PNA-positive acrosomal region. In *Lrrc37a*^−/−^ spermatozoa, this pattern was markedly disrupted, with irregular ATP1A1 distribution within the enlarged anterior head and loss of its characteristic spatial relationship with PNA (Fig. 6A, B). ATP1A4 likewise showed a defined localization pattern along the PNA-positive acrosomal region in control spermatozoa, whereas its localization became markedly disorganized in *Lrrc37a*^−/−^ spermatozoa and no longer maintained its characteristic spatial relationship with PNA (Fig. 6C, D). To determine when this localization defect emerged, ATP1A1 distribution was further analyzed during spermiogenesis. ATP1A1 localization was similar between *Lrrc37a*^+/+^ and *Lrrc37a*^−/−^ spermatids at steps 9–12, but became abnormal at steps 13–16 (Fig. 6E), coinciding with the onset of acrosomal abnormalities in *Lrrc37a*^−/−^ spermatids.

**Figure 6.**
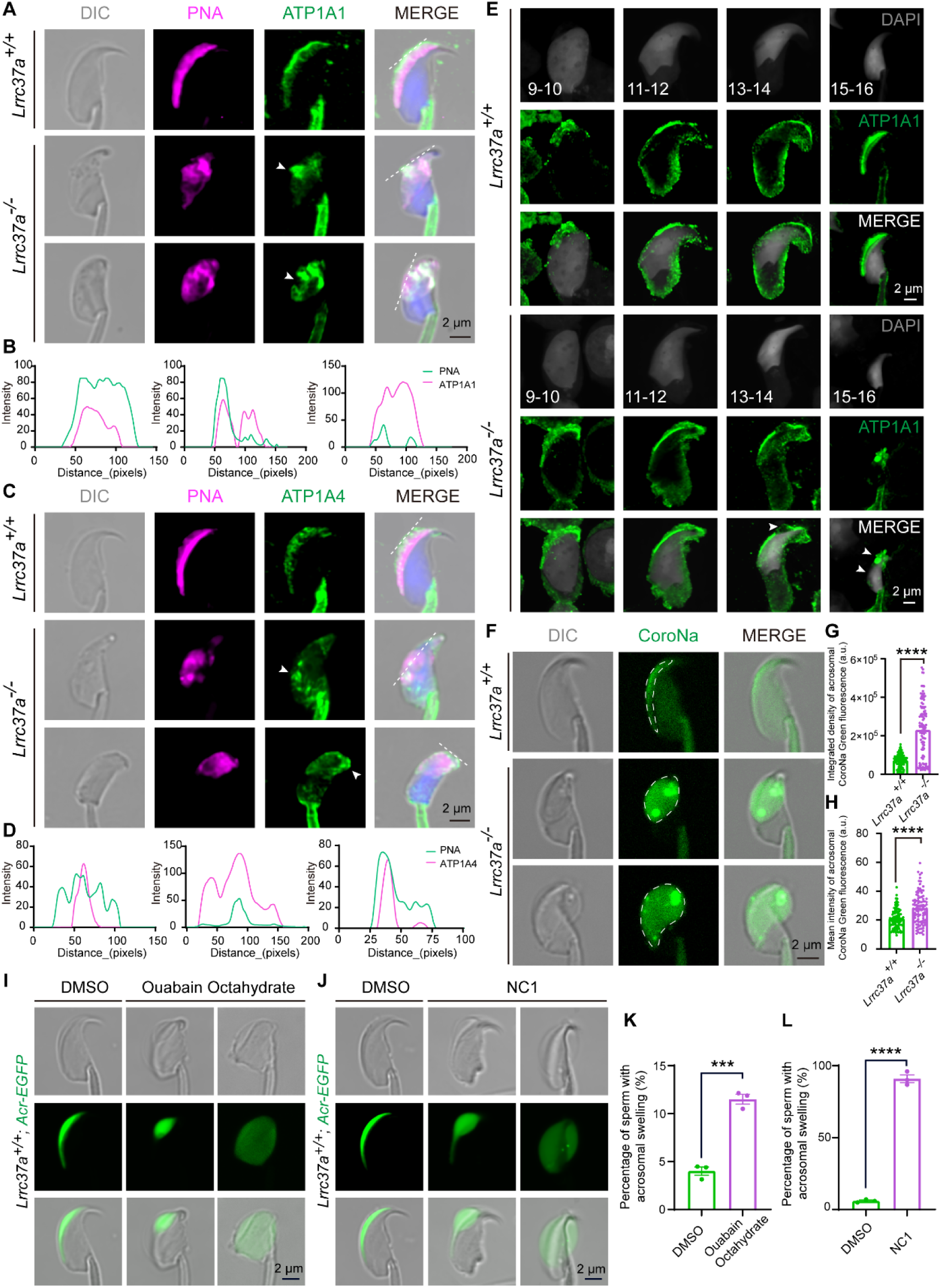
LRRC37A deficiency disrupts Na/K-ATPase localization and is accompanied by abnormal acrosomal Na⁺ accumulation. (A) Representative immunofluorescence images showing ATP1A1 and PNA in epididymal spermatozoa from *Lrrc37a*^+/+^ and *Lrrc37a*^−/−^ mice. White arrowheads indicate abnormal ATP1A1 localization. (B) Representative line-scan fluorescence intensity profiles of PNA and ATP1A1 along the dashed lines indicated in (A). (C) Representative immunofluorescence images showing ATP1A4 and PNA in epididymal spermatozoa from *Lrrc37a*^+/+^ and *Lrrc37a*^−/−^ mice. (D) Representative line-scan fluorescence intensity profiles of PNA and ATP1A4 along the dashed lines indicated in (C). (E) Immunofluorescence analysis of ATP1A1 in *Lrrc37a*^+/+^ and *Lrrc37a*^−/−^ spermatids during spermiogenesis. White arrowheads indicate abnormal ATP1A1 localization. DNA was counterstained with DAPI and is shown in gray. (F) Representative DIC and fluorescence images of epididymal spermatozoa from *Lrrc37a*^+/+^ and *Lrrc37a*^−/−^ mice stained with the Na⁺-sensitive fluorescent probe CoroNa Green. Dashed lines delineate the acrosomal region used for fluorescence quantification. (G, H) Quantification of the integrated density (G) and mean fluorescence intensity (H) of CoroNa Green within the acrosomal region of *Lrrc37a*^+/+^ and *Lrrc37a*^−/−^ spermatozoa. More than 100 spermatozoa in total from three mice per genotype were analyzed. Data are presented as the mean ± SEM. Two-tailed, unpaired Student’s t test: \*\*\*\**P* < 0.0001. (I) Live-cell analysis of acrosomal morphology in spermatozoa carrying the acrosomal EGFP reporter after treatment with DMSO or 1 mM ouabain octahydrate for 1 h. (J) Live-cell analysis of acrosomal morphology in spermatozoa carrying the acrosomal EGFP reporter after treatment with DMSO or 10⁻⁵ M Necrocide 1 (NC1) for 30 min. (K) Quantification of the percentage of spermatozoa exhibiting acrosomal swelling after treatment with DMSO or 1 mM ouabain octahydrate for 1 h. Three independent experiments were performed, with more than 200 spermatozoa analyzed in each experiment. Data are presented as the mean ± SEM. Two-tailed, unpaired Student’s t test: \*\*\**P* < 0.001. (L) Quantification of the percentage of spermatozoa exhibiting acrosomal swelling after treatment with DMSO or 10⁻⁵ M NC1 for 30 min. Three independent experiments were performed, with more than 200 spermatozoa analyzed in each experiment. Data are presented as the mean ± SEM. Two-tailed, unpaired Student’s t test: \*\*\*\**P* < 0.0001.

Given the established role of ATP1A1 and ATP1A4 in Na⁺ transport, Na⁺ distribution was assessed using the Na⁺-sensitive fluorescent probe CoroNa Green. *Lrrc37a*^−/−^ spermatozoa exhibited a marked accumulation of CoroNa Green fluorescence within the acrosomal compartment compared with *Lrrc37a*^+/+^ spermatozoa (Fig. 6F). Quantification of both integrated fluorescence density and mean fluorescence intensity confirmed significant increases in the *Lrrc37a*^−/−^ spermatozoa (Fig. 6G, H), demonstrating a marked disturbance of local Na⁺ homeostasis within the acrosomal region following loss of LRRC37A.

To assess whether perturbation of ion transport affects acrosomal morphology, spermatozoa carrying the acrosomal EGFP reporter were exposed to 1 mM ouabain for 1 h, a concentration reported to inhibit both NKAα4 and NKAα1 in mouse spermatozoa ^27^. Ouabain treatment significantly increased the proportion of spermatozoa exhibiting acrosomal swelling compared with vehicle-treated controls (Fig. 6I, K). Treatment with Necrocide 1 (NC1), which has been reported to induce cellular Na⁺ overload ^28^, produced a more pronounced phenotype, with marked enlargement of the acrosomal region in the majority of treated spermatozoa (Fig. 6J, L). Thus, disruption of Na/K-ATPase activity or cellular ion homeostasis was accompanied by acrosomal swelling, consistent with a functional relationship between local ion balance and acrosomal morphology.

## Discussion

Acrosome formation is initiated early in spermiogenesis through the delivery and fusion of Golgi-derived proacrosomal vesicles at the nuclear surface. Defects in this trafficking machinery therefore usually become apparent during the initial stages of acrosome assembly ^15^. For example, loss of GOPC causes acrosomal fragmentation in early round spermatids because Golgi-derived vesicles fail to fuse properly with the developing acrosome ^29^. PICK1 deficiency similarly disrupts proacrosomal vesicle trafficking and produces a globozoospermia-like phenotype ^30^. Germ-cell-specific deletion of *Atg7* also impairs proacrosomal vesicle trafficking and fusion, resulting in defective acrosome biogenesis ^31^. In contrast, acrosome formation in *Lrrc37a*^−/−^ spermatids remains largely intact through the Golgi, cap, and elongation phases, with pronounced enlargement becoming apparent only at steps 13–16. The defect therefore emerges after formation of the acrosomal compartment and is more pronounced in epididymal spermatozoa. This temporal pattern distinguishes LRRC37A deficiency from classical defects in acrosome biogenesis and indicates a requirement for LRRC37A during late acrosomal maturation rather than initial acrosome assembly.

The interaction of LRRC37A with ATP1A1 and ATP1A4 suggests that LRRC37A may contribute to ion regulation in the sperm head. Na/K-ATPase maintains transmembrane ion gradients in animal cells ^24^, and both the α1 and α4 catalytic isoforms are expressed in male germ cells and spermatozoa ^26^. *Atp1a4* knockout males are completely infertile, and their spermatozoa show severe motility defects and fail to fertilize wild-type oocytes ^25^. Super-resolution imaging has further shown that ATP1A1 and ATP1A4 are present in the sperm head as well as the flagellum and display distinct spatial distributions, with ATP1A4 undergoing redistribution during capacitation ^26^. In our study, ATP1A1 and ATP1A4 partially colocalized with LRRC37A in the sperm head, and both proteins showed marked changes in distribution following LRRC37A loss. ATP1A1 localization also became abnormal at steps 13–16, coinciding with the developmental onset of the acrosomal phenotype. These observations suggest that LRRC37A contributes to the proper localization of Na/K-ATPase isoforms in the sperm head, and that disruption of this organization accompanies the onset of acrosomal abnormalities.

Previous studies showed that *Atp1a4* knockout spermatozoa exhibit elevated intracellular Na⁺ at the whole-cell level ^25^. Recent work demonstrated that both ATP1A1 and ATP1A4 contribute to the sperm acrosome reaction and that ATP1A4-dependent Na⁺ regulation is important for a normal acrosomal response ^32^. Manipulation of extracellular Na⁺ or Na⁺ influx altered the abnormal acrosomal response of *Atp1a4* knockout spermatozoa, further linking Na⁺ homeostasis to acrosomal function ^32^. In *Lrrc37a*^−/−^ spermatozoa, Na⁺ was markedly increased within the acrosomal region, where pronounced swelling was also observed. Pharmacological perturbation of ion transport also induced abnormal acrosomal swelling. Ouabain treatment, which inhibits Na/K-ATPase activity, increased the proportion of spermatozoa exhibiting acrosomal swelling. NC1, which has been reported to induce cellular Na⁺ overload ^28^, also induced acrosomal swelling, but in a larger proportion of spermatozoa. The different responses to these treatments suggest that acrosomal swelling may be influenced by ion-regulatory processes beyond Na/K-ATPase activity. Pronounced acrosomal enlargement has not been reported as a characteristic feature of the *Atp1a4* knockout phenotype ^32^, further indicating that the phenotype of *Lrrc37a*^−/−^ spermatozoa cannot be explained by impaired ATP1A4 function alone. Together with the altered localization of ATP1A1 and ATP1A4 and the enrichment of transmembrane transport-related proteins in the LRRC37A interactome, these observations raise the possibility that LRRC37A-dependent ion homeostasis involves a broader set of ion-regulatory components. Whether other ion channels or transporters participate in LRRC37A-dependent acrosomal ion homeostasis remains to be determined.

Disturbance of ion homeostasis may alter the osmotic balance of the acrosomal compartment, promoting water influx and compartmental swelling. Spermatozoa are particularly sensitive to osmotic changes, and defective volume regulation can result in excessive swelling and structural deformation. The acrosome itself has also been shown to exhibit distinct osmotic behavior, with changes in tonicity affecting acrosomal volume and shape ^2,12,13^. The increased Na⁺ detected in the acrosomal region of *Lrrc37a*^−/−^ spermatozoa may therefore promote water entry and contribute to the pronounced acrosomal swelling. Although water movement across the acrosomal membranes was not directly measured, the induction of acrosomal swelling following pharmacological perturbation of ion homeostasis supports this possibility.

This requirement for ion and volume regulation may become particularly important as spermatozoa mature and encounter changing extracellular environments. During transit through the male and female reproductive tracts, sperm are exposed to substantial changes in osmolality and ionic composition and must maintain appropriate volume regulation ^2,12^. The appearance of acrosomal swelling during late spermiogenesis and its greater prominence in epididymal spermatozoa may therefore reflect an increasing requirement for maintaining the ionic and osmotic stability of the acrosomal compartment during sperm maturation.

The membrane abnormalities observed in *Lrrc37a*^−/−^ spermatozoa occurred in parallel with disruption of local ion homeostasis. The plasma membrane, outer acrosomal membrane, and inner acrosomal membrane became irregular, accompanied by expansion of the acrosomal compartment, reduced electron density, abnormal vesicular structures, and altered PNA and SPACA1 distributions. LRRC37A itself was positioned toward the outer acrosomal region, where it showed extensive overlap with PNA and lay external to SPACA1. Ionic and osmotic imbalance can influence compartment volume and membrane morphology, raising the possibility that at least part of the structural phenotype develops secondary to disturbed acrosomal ion homeostasis. Such abnormalities would be expected to impair the functional competence of the acrosomal region. Consistent with this, *Lrrc37a*^−/−^ spermatozoa showed reduced zona pellucida binding and a decreased acrosome reaction, together with complete failure of fertilization.

Recent reports have also implicated LRRC37A in acrosomal integrity and sperm-head morphogenesis ^33^, including a recent preprint proposing a role in the organization of the inner acrosomal membrane–perinuclear theca–nuclear envelope complex ^34^. These studies have largely focused on the structural role of LRRC37A in the acrosome and acrosome–nuclear interface, without addressing its relationship with ion-transport proteins or local ionic homeostasis in the sperm head. The present study was conceived and conducted independently of these reports and extends current understanding of LRRC37A by linking it to acrosomal ion homeostasis. Collectively, our study demonstrates that LRRC37A is required for acrosomal ion homeostasis and male fertility. Loss of LRRC37A disrupts ATP1A1 and ATP1A4 localization and is accompanied by increased Na⁺-sensitive fluorescence in the acrosomal region, pronounced acrosomal swelling, and complete fertilization failure. These findings link LRRC37A-dependent ion regulation to the maintenance of normal acrosomal physiology. The evolutionary conservation of LRRC37A across internally fertilizing vertebrates and its presence in human spermatozoa suggest that this function may be conserved and relevant to human male fertility.

## Experimental procedures

### Animals

The mouse *Lrrc37a* gene (ENSMUST00000153273.3) is located on chromosome 11 with 14 exons, a portion of the coding region of Exon 1 was selected as the CRISPR-Cas9 target region. *Lrrc37a* knockout mice were generated by Rise Mice Biotechnology Co., Ltd (Zhaoqing, China). *Lrrc37a*-HA knock-in mice were generated by Cyagen Biosciences using the CRISPR-Cas9 system. For *Lrrc37a*-HA line, a gRNA targeting *Lrrc37a*, a donor vector containing “HA tag” cassette, and Cas9 mRNA were co-injected into fertilized eggs. F0 founders were screened by PCR and sequencing, followed by breeding with wild-type mice to confirm germline transmission and obtain F1 offspring. B6D2-Tg (CAG/Su9-DsRed2, Acr3-EGFP) RBGS002Osb mice (no. 03743), which express DsRed2 in sperm mitochondria and EGFP in the acrosome ^19^, were used for live imaging of sperm acrosomal morphology.

Primer sequences used for genotyping are listed below.

For knockout genotyping,

*Lrrc37a* F: CACTTGCTAAGGAGGTTGTCG

*Lrrc37a* R1: TCACTTAGGACCTCTTGTTGGACT

Wildtype allele: 673 bp

*Lrrc37a* F: CACTTGCTAAGGAGGTTGTCG

*Lrrc37a* R2: GTCTGAGTCTCTTCCTGAACTGG

Mutant allele: 389 bp For knockin genotyping,

F4: 5’-TGATCCAACATGCCCAACTAGCTATG-3’

R3: 5’-TCTAGGAACTTTTGTTCCACAAGGC-3’

Wildtype allele: 356 bp

F3: 5’-CGATGTTCCAGATTACGCTTG-3’

R1: 5’-TAGCTTCTCTAGGCACCGGATC-3’

Mutant allele: 391 bp

All mice were maintained under specific pathogen-free conditions at 23 ± 2 °C with a reverse light cycle and free access to food and water. Ethical approval for all animal experimental procedures was provided by the Laboratory Animal Welfare and Ethics Committee of Guangzhou Women and Children’s Medical Center (Ethics No. KTDW-2024-00337).

### Human sperm sample preparation

Semen samples were collected from healthy young Chinese volunteers who met the study inclusion criteria following medical screening. Samples were purified using a 40% PureSperm density gradient and washed with PBS before immunofluorescence staining. All participants provided written informed consent. The study was approved by the Medical Ethics Committee of Guangzhou Women and Children’s Medical Center (approval no. 2024126A01) and conducted in accordance with the Declaration of Helsinki.

### Plasmids

Mouse *Atp1a1* and *Atp1a4* were amplified from mouse testis cDNA and cloned into the pCMV-HA vector using the Clone Express Ultra One Step Cloning Kit (C115, Vazyme). The mouse LRRC37A-FLAG expression plasmid was constructed by Beijing SYKM Gene Biotechnology Co., Ltd.

### Cell culture and transfection

HEK293T cells (Cat# GNHu17; National Collection of Authenticated Cell Cultures, Chinese Academy of Sciences) were cultured in DMEM (Cat# C11995500BT, Gibco) containing 10% fetal bovine serum (Cat# 16000044, Gibco) and 1% penicillin–streptomycin (Cat# 15140122, Gibco). Cells were maintained at 37°C in a humidified atmosphere containing 5% CO₂ and routinely passaged with trypsin–EDTA (Cat# 25200072, Gibco). Transfection was performed when cultures reached approximately 70–80% confluence using LipoMax DNA transfection reagent (Cat# 32110, Sudgen), following the manufacturer’s protocol.

### Antibodies

Primary antibodies used in this study were rabbit anti-LRRC37A (homemade, Dia-an Biotech, 1:1000 for WB, 1:50 for IF), rabbit anti ATP1A4 (homemade, Dia-an Biotech, 1:50 for IF), rabbit anti-ATP1A1 (14418-1-AP, 1:50 for IF), rabbit anti-SPACA1 (ab191843, Abcam, 1:100 for IF), mouse anti-SP56 (55101, QED Bioscience, 1:100 for IF), mouse anti-α-Tubulin (AC012, Abclonal, 1:3000 for WB, 1:100 for IF), rabbit anti-FLAG (PM020, MBL, 1:1000 for WB), mouse anti-HA (2367S, Cell Signaling Technology, 1:2000 for WB, 1:100 for IF). Secondary antibodies used in this study were goat anti-mouse FITC (ZF-0312, Zhong Shan Jin Qiao, 1:200), goat anti-mouse TRITC (ZF0313, Zhong Shan Jin Qiao, 1:200), goat anti-rabbit FITC (ZF-0311, Zhong Shan Jin Qiao, 1:200), goat anti-rabbit TRITC (ZF-0316, Zhong Shan Jin Qiao, 1:200), Alexa Fluor 594-conjugated PNA (Thermo Fisher Scientific, Cat. No. L32459, 1:400) and TRITC-conjugated phalloidin (YEASEN, Cat. No. 40734ES75, 1:1000).

### Assessment of fertility

Fertility was evaluated in two-month-old males of each genotype by continuous mating with two 7–8-week-old wild-type C57BL/6 females. Vaginal plugs were examined daily, and plug-positive females were separated and monitored for pregnancy and parturition. Delivery dates and litter sizes were recorded throughout the study. Reproductive performance was assessed over a 6-month period based on pregnancy outcomes and litter production.

### Reverse transcription-quantitative PCR (RT-PCR)

Total RNA was isolated from various tissues of 8-week-old wild-type (WT) mice using the FastPure Cell/Tissue Total RNA Isolation Kit V2 (Vazyme, RC112-01). RNA concentrations were determined using a NanoDrop spectrophotometer. First-strand complementary DNA (cDNA) was generated using 5× PrimeScript RT Master Mix (Takara, RR036A) following the manufacturer’s protocol. The synthesized cDNA served as the template for subsequent PCR amplification. The following primer sequences were used:

*β-actin* forward: 5′-GGCTGTATTCCCCTCCATCG-3′;

*β-actin* reverse: 5′-CCAGTTGGTAACAATGCCATGT-3′;

*Lrrc37a* forward: 5′-ACCAAATGTCTCCACAAGCAC-3′;

*Lrrc37a* reverse: 5′-GGCGATTTTCGCTGAGAATTAGTTT-3′.

### Immunoprecipitation

Testes were homogenized in lysis buffer (Beyotime, P0013) supplemented with protease inhibitor cocktail (Roche, 04693132001). Lysates were briefly sonicated and incubated on ice for 30 min, followed by centrifugation at 12,000 rpm for 20 min at 4°C. The supernatants were collected, and equal amounts of protein were incubated with the indicated primary antibodies overnight at 4°C with gentle rotation. Protein A-conjugated beads were then added and incubated for 4 h at 4°C. The beads were washed three times with ice-cold lysis buffer, and bound proteins were eluted for subsequent immunoblotting or LC–MS/MS analysis.

### Proteomics analysis

Tryptic peptides were dissolved in solvent A (0.1% formic acid in water) and loaded onto a homemade reversed-phase analytical column (15 cm length, 100 μm i.d.). Peptide separation was performed on a Vanquish Neo UPLC system (Thermo Fisher Scientific) at a flow rate of 600 nL/min using solvent B containing 0.1% formic acid in 80% acetonitrile. The gradient was as follows: 4% B for 0–0.5 min, 4–7.5% B for 0.5–0.6 min, 7.5–30% B for 0.6–8.1 min, 30–44% B for 8.1–9.6 min, 44–55% B for 9.6–10 min, 55–99% B for 10–10.5 min, and 99% B for 10.5–12 min. Peptides were analyzed using a timsTOF HT mass spectrometer operated in data-independent parallel accumulation–serial fragmentation (dia-PASEF) mode with an electrospray voltage of 1.6 kV. Full MS scans were acquired over an *m/z* range of 300–1500, with 12 PASEF MS/MS scans acquired per cycle. The MS/MS scan range was 395–1080 *m/z*, with an isolation window of 15 *m/z*. DIA data were processed using DIA-NN version 1.8. Spectra were searched against the *Mus musculus* database (Mus_musculus_10090_UP_20241025_seqkit.fasta; 84,193 entries) concatenated with a reverse decoy database. Trypsin/P was specified as the cleavage enzyme, allowing up to one missed cleavage. N-terminal methionine excision and carbamidomethylation of cysteine were specified as fixed modifications. The false discovery rate was controlled at <1%.

### Phylogenetic analysis

Amino acid sequences of LRRC37A from 10 species were downloaded from the NCBI (https://www.ncbi.nlm.nih.gov/protein) and UniProt databases (http://www.uniprot.org). A phylogenetic tree was constructed using the neighbor-joining (NJ) method in MEGA 11.0. Evolutionary distances were calculated using the p-distance method, and branch support was assessed by bootstrap analysis with 1,000 replicates.

### Immunoblotting

Tissues were homogenized in lysis buffer (Beyotime, P0013) containing protease inhibitor cocktail (Roche, 04693132001) and 1 mM phenylmethylsulfonyl fluoride (PMSF; Amresco, 0754). After incubation on ice for 30 min, lysates were clarified by centrifugation at 12,000 rpm for 20 min at 4°C, and the supernatants were collected.

Proteins were separated by SDS–PAGE and transferred to nitrocellulose membranes. After blocking with 5% skim milk in PBS for 1 h at room temperature, membranes were incubated with primary antibodies overnight at 4°C. Following PBS washes, HRP-conjugated secondary antibodies were applied for 1 h at room temperature. Chemiluminescent signals were visualized using a Touch Imager Pro system (e-BLOT).

### Sperm motility assessment

The cauda epididymides were isolated from males of each genotype, and spermatozoa were released into Human Tubal Fluid medium (Sigma-Aldrich, catalog no. MR-070-D) at 37 °C for 10 min. The swim-up fraction was examined using a 10× phase objective on an Axiolab 5 microscope (Carl Zeiss Microscopy, LLC, White Plains, NY, USA) equipped with a CCD camera (Hamilton Thorne, Beverly, MA, USA). Total and progressive motility were quantified by computer-assisted semen analysis (CASA) using CEROS software, and sperm concentration was determined with a hemocytometer.

### Immunofluorescence

For sperm immunofluorescence, epididymal spermatozoa were released into PBS at 37°C for 10 min and spread onto glass slides. After air-drying, samples were fixed with 4% paraformaldehyde for 5 min, washed with PBS, and blocked with 5% bovine serum albumin (BSA). Samples were incubated with primary antibodies overnight at 4°C, followed by the appropriate secondary antibodies and DAPI counterstaining. Images were acquired using a Nikon AXR microscope (Tokyo, Japan) or a Multimodality Structured Illumination Microscopy system (NanoInsights-Tech Co., Ltd.). Colocalization analysis was performed using ImageJ software.

### Acrosomal sodium imaging

Cauda epididymal spermatozoa were released into modified Tyrode’s medium containing 135 mM NaCl, 5.0 mM KCl, 1.2 mM KH₂PO₄, 1.2 mM MgSO₄, 5.5 mM glucose, 0.8 mM pyruvate, 4.8 mM lactate, and 20 mM HEPES (pH 7.4). CoroNa Green, AM (C36676, Thermo Fisher Scientific) was premixed with Pluronic F-127 (P6866, Invitrogen) and added to the sperm suspension at a final concentration of 10 μM. Spermatozoa were incubated for 30 min at 37°C, washed with modified Tyrode’s medium, and subjected to confocal fluorescence imaging. CoroNa Green fluorescence (Ex/Em = 492/516 nm) in the acrosomal region was quantified using ImageJ.

### Live-cell analysis of acrosomal morphology following disruption of ion homeostasis

Cauda epididymal spermatozoa were released into the same modified Tyrode’s medium and treated with 1 mM ouabain octahydrate (MedChemExpress, Cat. No. HY-B0542) for 1 h or 10 μM Necrocide-1 (NC1; MedChemExpress, Cat. No. HY-14307) for 30 min at 37°C. Fluorescence and corresponding bright-field images were acquired and overlaid to evaluate acrosomal morphology in living spermatozoa. The percentage of spermatozoa exhibiting abnormal acrosomal swelling was quantified.

### Transmission electron microscopy

For epididymal samples, caput, corpus, and cauda epididymides were fixed overnight at 4°C in 2.5% glutaraldehyde prepared in 0.1 M cacodylate buffer. For testicular samples, testes were dissected and fixed under the same conditions. After washing, tissues were cut into approximately 1 mm³ pieces, post-fixed in 1% osmium tetroxide (OsO₄) for 1 h, dehydrated through a graded acetone series, and embedded in resin. Ultrathin sections were prepared, stained with uranyl acetate and lead citrate, and examined using a JEOL JEM-1400 transmission electron microscope (JEOL Ltd., Akishima, Tokyo, Japan).

### Scanning electron microscopy

Spermatozoa were separately collected from the caput, corpus, and cauda epididymides and released into PBS at 37°C for 10 min. After centrifugation at 500 × *g* for 5 min, samples were washed with phosphate buffer and fixed in 2.5% glutaraldehyde overnight. The specimens were then dehydrated through a graded ethanol series, dried, sputter-coated with gold, and examined using a Hitachi SU8010 scanning electron microscope (Hitachi High-Tech Corporation, Tokyo, Japan).

### Superovulation and *in vitro* fertilization

For superovulation, female mice were injected intraperitoneally with 5 IU of pregnant mare serum gonadotropin (PMSG; 110,254,564, Ningbo Second Hormone Company), followed 46– 48 h later by 5 IU of human chorionic gonadotropin (hCG; 110,251,281, Ningbo Second Hormone Company). At 13 h post-hCG, cumulus-oocyte complexes (COCs) were surgically collected from the ampulla of the oviducts.

Cauda epididymal spermatozoa from sexually mature *Lrrc37a*^+/+^ and *Lrrc37a*^−/−^ males were preincubated in human tubal fluid (HTF) medium (Sudgen Biotechnology, China) for 1 h at 37 °C under 5% CO₂ to induce capacitation. COCs were treated with 1 mg/mL hyaluronidase to disperse cumulus cells, and Tyrode’s solution (T1788, Sigma-Aldrich) was used to remove the zona pellucida (ZP). Cumulus-intact, cumulus-free and ZP-free oocytes were then incubated with spermatozoa at a final concentration of 4 × 10⁵/mL in 100 µL drops of HTF medium. After 3 h of co-incubation, fertilized oocytes were washed in fresh HTF drops to remove abnormal oocytes, and all manipulations were performed at 37 °C on a TPiE-SMZR constant-temperature plate (TOKAI HIT). Fertilization success was assessed at 6 h post-insemination by the presence of two distinct pronuclei.

### Sperm ZP binding assay

Capacitated spermatozoa (4 × 10⁵/mL) were incubated with ovulated oocytes in 100 µL drops of HTF medium for 1 h at 37 °C in a humidified atmosphere of 5% CO₂. After incubation, samples were fixed in 4% paraformaldehyde (PFA) containing 0.5% Triton X-100 for 10 min, followed by staining with 4′,6-diamidino-2-phenylindole (DAPI) to visualize sperm nuclei. The number of spermatozoa bound to the zona pellucida was quantified from z-projections acquired using a confocal microscope.

### Oocyte and embryo immunofluorescence staining

For immunofluorescence staining, oocytes or embryos were fixed and permeabilized simultaneously in 4% paraformaldehyde (PFA) containing 0.5% Triton X-100 for 10 min at room temperature. After blocking with 1% bovine serum albumin (BSA; AP0027, Amresco, USA) in PBS for 30 min, samples were incubated with Phalloidin for 30 min at 37 °C, followed by DAPI staining for 5 min to visualize nuclei. Finally, stained oocytes and embryos were mounted onto glass slides and imaged using a Nikon AXR confocal microscope (Tokyo, Japan).

### Statistical analysis

All experiments were performed at least three independent times, and data are presented as mean ± SEM. Statistical comparisons between two groups were performed using unpaired, two-tailed Student’s *t*-tests. *P* < 0.05 was considered statistically significant, with significance levels indicated as follows: *P <* 0.01 (**)*, P* < 0.001 (***), and *P* < 0.0001 (****).

## Data availability

All data needed to evaluate the conclusions in the paper are present in the paper and/or the Supplementary Materials.

## Acknowledgements

We thank Dr. Masaru Okabe (Osaka University) for providing the B6D2-Tg (CAG/Su9-DsRed2, Acr3-EGFP) RBGS002Osb transgenic mice.

## Author contributions

L., and B. W. Conceptualization; W. L., B. W., L. W., J. L., and T. T. Methodology; B. W., L. W., J. L., and C. L. Investigation; B. W., L. W., and J. L. Validation; B. W., L. W., J. L., C. L., Y. M., T. H., X. H., and Q. S. Visualization; B. W., L. W., and J. L. Data curation; B. W., L. W., and J. L. Writing-original draft; W. L. and B. W. Writing-review and editing. W. L., B. W., L. W., and T. T. Funding acquisition; W. L. and B. W. Project administration; All authors assisted in data collection, interpreted the data, provided critical input to the manuscript, and approved the final manuscript.

## Funding and additional information

This work was supported by the National Natural Science Foundation of China (Grant No. 32500759, 32230029, 32400709, 32500060), and the China Postdoctoral Science Foundation (2025M772781, 2024M760638, 2024M760634, 2026T190696), and the Postdoctoral Fellowship Program of CPSF (GZC20251824).

## Abbreviations and nomenclature

LRRC37A: Leucine-rich repeat-containing protein 37A
ATP1A1: Na/K-ATPase α1 subunit
ATP1A4: Na/K-ATPase α4 subunit
CASA: Computer-assisted sperm analysis
ZP: Zona pellucida
NC1: Necrocide-1
COC: Cumulus–oocyte complex
HTF: Human tubal fluid
H&E: Hematoxylin and eosin
PAS: Periodic acid-Schiff
SEM: Scanning electron microscopy
TEM: Transmission electron microscopy

## Supplementary Materials

**Figure S1.**
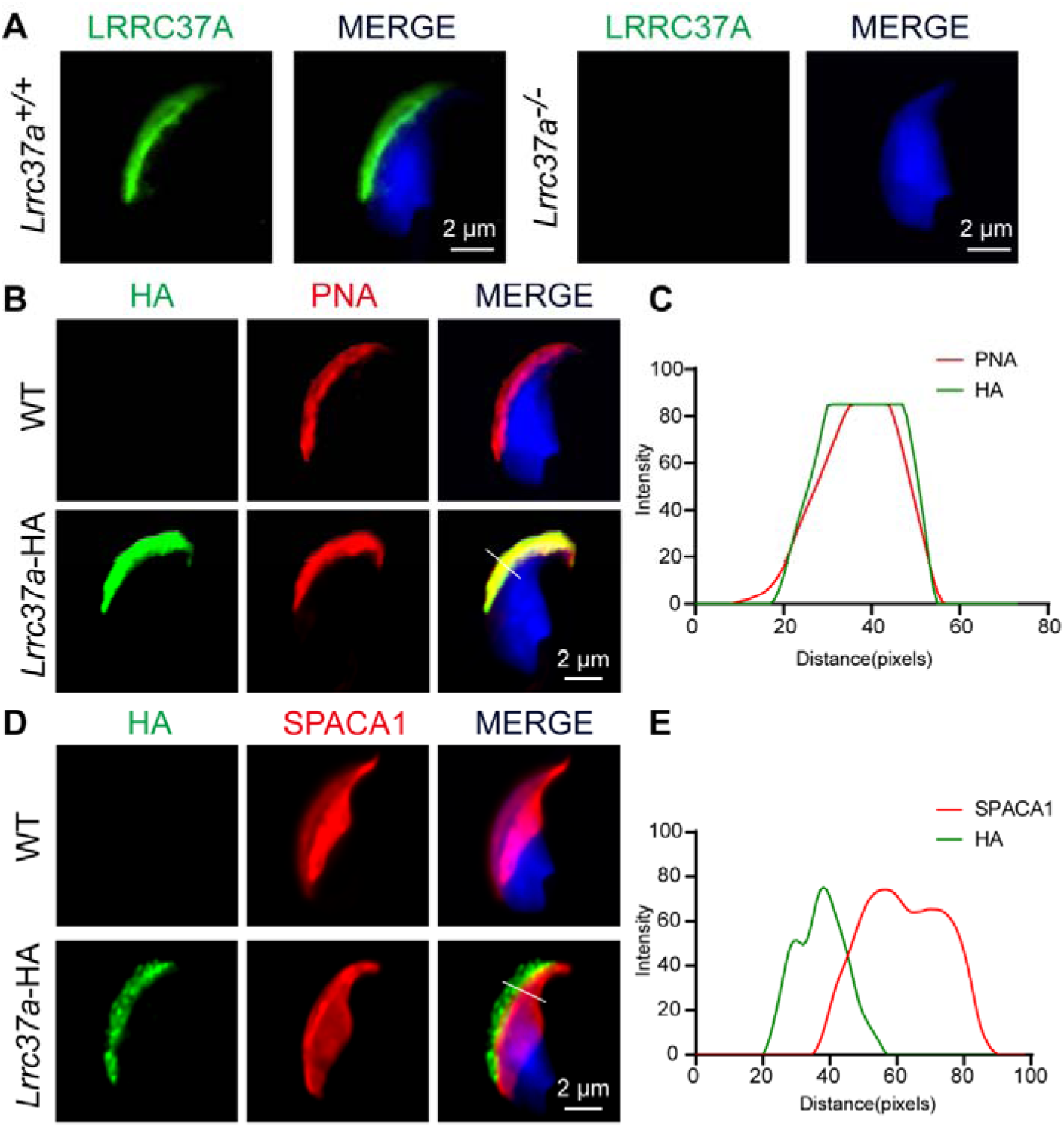
Validation of LRRC37A deficiency and HA-tagged LRRC37A localization in spermatozoa. (A) Immunofluorescence analysis of LRRC37A in mature spermatozoa from *Lrrc37a*^+/+^ and *Lrrc37a*^−/−^ mice. LRRC37A signal in the sperm head is absent in *Lrrc37a*^−/−^ spermatozoa. (B, C) Confocal immunofluorescence images showing the spatial relationship between HA-tagged LRRC37A and PNA in mature spermatozoa, with corresponding fluorescence intensity profiles. (D, E) Confocal immunofluorescence images showing the spatial relationship between HA-tagged LRRC37A and SPACA1 in mature spermatozoa, with corresponding fluorescence intensity profiles.

**Table S1 (separate file). Proteins identified in LRRC37A-HA and wild-type control immunoprecipitates from mouse testis.**

